# Optimized Multiple Amplification Protocol for the Production of Allogeneic Human Vγ9Vδ2 T Lymphocytes for Adoptive Cell Transfer Immunotherapy

**DOI:** 10.64898/2026.03.25.714172

**Authors:** Noémie Joalland, Laura Lafrance, Emmanuel Scotet

## Abstract

Cancer remains a major therapeutic challenge despite substantial advances in diagnosis and treatment, including immune checkpoint blockade. Among emerging immunotherapeutic approaches, adoptive cell transfer (ACT) has attracted growing interest. Human peripheral Vγ9Vδ2 T cells are promising candidates for ACT because they combine rapid and potent antitumor functions with major histocompatibility complex (MHC)-independent tumor recognition, enabling allogeneic use with limited risk of graft-versus-host disease. This raises the possibility of generating standardized Vγ9Vδ2 T-cell banks from healthy donors for off-the-shelf immunotherapy. Here, we provide preclinical evidence supporting the suitability of allogeneic human Vγ9Vδ2 T cells for ACT. We characterized peripheral blood Vγ9Vδ2 T cells from healthy donors after successive antigen-specific and non-specific amplification steps, assessing their phenotype, effector functions, and metabolic state. Amplified cells maintained a strong pro-inflammatory Th1-like profile, preserved cytotoxic activity, and did not produce immunoregulatory cytokines. They also displayed high purity, a predominant effector memory phenotype, reduced expression of several inhibitory immune checkpoints, and sustained antitumor reactivity. Altogether, these findings support the development of allogeneic Vγ9Vδ2 T-cell products as a scalable platform for next-generation cancer immunotherapies.

## INTRODUCTION

Cancer remains one of the leading causes of mortality worldwide despite major advances in diagnosis and treatment, including surgery, chemotherapy, targeted therapies, and immune checkpoint blockade. In this context, immunotherapy has emerged as a major therapeutic breakthrough by harnessing the immune system to recognize and eliminate malignant cells. Among immunotherapeutic approaches, adoptive cell transfer (ACT) has shown considerable promise for inducing potent and durable antitumor responses (Rosenberg & Restifo, 2015). ACT relies on the isolation, *ex vivo* expansion and/or modification, and reinfusion of immune effector cells into patients in order to enhance anticancer immunity. This strategy can partially overcome tumor immune evasion by directly providing activated immune cells with predefined antitumor properties. While most ACT approaches have focused on CD8+ and CD4+ αβ T cells or natural killer (NK) cells, γδ T cells have progressively emerged as attractive alternative effectors for cellular immunotherapy (Silva-Santos et al., 2015).

γδ T cells occupy a singular position at the interface between innate and adaptive immunity. They combine rapid effector responsiveness with cytotoxic and cytokine-producing capacities, while recognizing stressed or transformed cells in a largely major histocompatibility complex (MHC)-independent manner (Vantourout & Hayday, 2013; Chien et al., 2014). This property is particularly relevant in cancer, as many tumors escape αβ T-cell surveillance through MHC downregulation. Human γδ T cells are classically subdivided according to their δ-chain usage. Vδ2 negative γδ T cells are mainly tissue-resident and recognize a broader range of ligands, including stress-associated molecules and CD1d-presented lipids (Zhao et al., 2018), whereas Vδ2 positive cells, most commonly paired with the Vγ9 chain in peripheral blood, respond to phosphoantigens (PAgs) such as isopentenyl pyrophosphate (IPP) and hydroxymethylbutenyl pyrophosphate (HMBPP), metabolites of the isoprenoid pathway (Tanaka et al., 1995). Their activation depends on butyrophilin family members involved in phosphoantigen sensing, notably BTN3A1 and BTN2A1 (Harly et al., 2012; Cano et al., 2021).

Among human γδ T-cell subsets, Vγ9Vδ2 T cells are the best characterized in the context of cancer immunotherapy. They are abundant in peripheral blood, can be selectively activated with phosphoantigens or aminobisphosphonates such as zoledronate, and display pleiotropic antitumor functions, including direct cytotoxicity, secretion of pro-inflammatory cytokines such as IFN-γ and TNF-α, and crosstalk with other immune populations (Dunne et al., 2010). Their low alloreactivity and lack of graft-versus-host disease (GvHD) induction make them particularly appealing for allogeneic ACT strategies (Dieli et al., 2007). These features have supported the development of clinical approaches based on *in vivo* stimulation or *ex vivo* expansion of Vγ9Vδ2 T cells, with encouraging but still heterogeneous outcomes (Wilhelm et al., 2003; Bennouna et al., 2008; Fournié et al., 2013).

However, several limitations still hamper the broader clinical use of Vγ9Vδ2 T cells. As for other ACT products, therapeutic efficacy depends on the ability to generate sufficient cell numbers, preserve functional competence during ex vivo manipulation, and ensure persistence after transfer (Rosenberg & Restifo, 2015; Zhang et al., 2023). In addition, the immunosuppressive tumor microenvironment can constrain the activity of transferred cells, while repeated stimulation during manufacturing may alter their phenotype, differentiation state, metabolic fitness, and proliferative potential. Although most studies have relied on a single round of specific expansion before administration, the production of standardized allogeneic cell banks will likely require more scalable manufacturing strategies capable of generating large quantities of highly pure cells without compromising their antitumor properties.

In this study, we investigated whether successive rounds of specific and non-specific amplification could provide such a strategy for the generation of polyclonal Vγ9Vδ2 T-cell products. Starting from peripheral blood mononuclear cells of healthy donors, Vγ9Vδ2 T cells were first specifically expanded using BrHPP or zoledronate, and then subjected to repeated non-specific restimulation with PHA and feeder cells. We compared cells at different stages of the amplification process in terms of cytokine profile, purity, differentiation status, inhibitory checkpoint expression, tumor reactivity, metabolic activity, and long-term proliferative potential. Our results show that repeated amplification generates highly pure Vγ9Vδ2 T-cell populations with preserved or enhanced effector properties, reduced expression of several inhibitory immune checkpoints, and marked metabolic remodeling. Altogether, these findings support the feasibility of producing large numbers of functional Vγ9Vδ2 T cells for the establishment of allogeneic cell banks and further strengthen the rationale for their development in adoptive immunotherapy.

## MATERIAL AND METHODS

### Sorting of γδ T Lymphocytes

Human γδ T lymphocytes were isolated from peripheral blood mononuclear cells (PBMCs) obtained from healthy donor blood samples provided by the Etablissement Français du Sang (EFS, Nantes, France) using Ficoll density centrifugation (Eurobio, Les Ulis, France). Untouched γδ T cells were sorted using the Human Gamma/Delta T Cell Isolation Kit (Stemcell Technologies, Vancouver, Canada; #19255) according to the manufacturer’s instructions. Briefly, PBMCs were resuspended in PBS containing 2% FBS and 1 mM EDTA at a concentration of 5 × 10^7 cells/mL. The isolation cocktail was added, followed by a 15-minute incubation. Magnetic particles were then added, and the mixture was incubated for an additional 10 minutes. The tube was placed in a magnet, and the supernatant containing γδ T cells was recovered after 5 minutes. This process was repeated twice to ensure high purity. The purity of the isolated cells was verified by flow cytometry.

### Amplification of Vγ9Vδ2 T Lymphocytes

For the specific expansion of human Vγ9Vδ2 T lymphocytes, PBMCs were incubated with 3 µM bromohydrin pyrophosphate (BrHPP; Innate Pharma, Marseille, France) or 5 µM zoledronic acid (Sigma Aldrich, Saint-Louis, MI, USA) in RPMI medium supplemented with 10% heat-inactivated FBS, 2 mM L-glutamine, 10 mg/mL streptomycin, 100 IU/mL penicillin (all from Gibco, Thermo Fisher, Waltham, MA, USA), and 100 IU/mL human recombinant IL-2 (Proleukin, Novartis, Basel, Switzerland). After 4 days, the culture medium was supplemented with 300 IU/mL IL-2. On day 21, the purity of the cell culture was assessed by flow cytometry, and only preparations with a purity > 80% were used. These pure human Vγ9Vδ2 T lymphocytes were then non-specifically expanded using mixed feeder cells composed of 35 Gy-irradiated Epstein-Barr Virus-transformed human B lymphocytes and PBMCs (each from 3 different donors) and PHA-L (Sigma) in RPMI medium supplemented with 10% heat-inactivated fetal calf serum, 2 mM L-glutamine, 10 mg/mL streptomycin, 100 IU/mL penicillin (all from Gibco), and 300 IU/mL recombinant human IL-2 (Novartis). After three weeks of culture and purity verification, Vγ9Vδ2 T cells could be used and/or amplified again using the same protocol. On day 21, Vγ9Vδ2 T cells were counted, and the number of divisions was calculated as follows: ln(number of cells at day 21/number of cells at day 0)/ln(2).

### Flow Cytometry

After sorting and/or amplification, human γδ T lymphocytes were stained with PC5-labeled anti-human pan-γδ mAb (#IMMU510, Beckman Coulter, Brea, CA, USA) and FITC-labeled anti-human TCR Vδ2 mAb (#IMMU389, Beckman Coulter) to assess population purity. The differentiation state of γδ T cells was analyzed using PE-labeled anti-human CD27 mAb (1A4CD27, Beckman Coulter) and APC-labeled anti-human CD45RA mAb (HI100, BD Biosciences, Franklin Lakes, NJ). The expression of inhibitory immune checkpoints was evaluated by staining with PE-labeled anti-human PD-1 mAb (329905, Biolegend, San Diego, CA), BV421-labeled anti-human CTLA-4 mAb (369606, Biolegend), AF488-labeled anti-human LAG-3 mAb (369325, Biolegend), and AF647-labeled anti-human Tim3 mAb (565558, BD Biosciences). Data were acquired using a FACSCalibur, Accuri C6 PLUS, or Canto II flow cytometer (all from BD Biosciences, Franklin Lakes, NJ, USA) and analyzed using Accuri software (BD Biosciences) or FlowJo software (Treestar, Ashland, OR, USA).

### Cytokine Quantification and Intracellular Staining

Human γδ T cells were plated at 1 × 10^5 cells per well in 100 µL of RPMI containing 10% FBS and activated with 30 µM BrHPP (Innate Pharma) or a mix of 500 ng/mL PMA and 1 µM ionomycin (both from Sigma). For cytokine quantification, supernatants were collected 6 hours after activation and analyzed using the LEGENDplex Human Th Cytokine Panel (#740001, Biolegend) according to the manufacturer’s instructions. This kit allows the quantification of 13 cytokines: IL-5, IL-13, IL-2, IL-6, IL-9, IL-10, IFN-γ, TNF-α, IL-17A, IL-17F, IL-4, IL-21, and IL-22. A standard curve was generated to quantify cytokine concentrations. For intracellular staining, γδ T cells were activated in the presence of 10 µM monensin (Sigma). After 6 hours, cells were fixed in 4% PFA, permeabilized, and stained with PE-labeled anti-human IFN-γ mAb (4S.B3, eBiosciences). Staining was analyzed by flow cytometry.

### CD107a Reactivity Assays

For the CD107a surface mobilization assay, PC3 prostate cancer cells or RAJI lymphoma cells were sensitized overnight with different concentrations of zoledronic acid (Sigma Aldrich). The co-culture with allogeneic human γδ T lymphocytes was performed at an effector-to-target ratio of 1:1 in RPMI medium containing 5 µM monensin (Sigma) and Alexa Fluor 647-labeled anti-human CD107a mAb (#H4A3, Biolegend). After 4 hours of incubation, Vγ9Vδ2 T lymphocytes were collected, stained with FITC-labeled anti-human TCR Vδ2 mAb (#IMMU389, Beckman Coulter), and analyzed by flow cytometry.

### Metabolic Analysis Using Seahorse

Mitochondrial oxygen consumption rate (OCR) and extracellular acidification rate (ECAR) were measured in medium buffered to pH 7.4 with NaOH and supplemented with glucose (25 mM), pyruvate (1 mM), glutamine (2 mM), and 1% FBS using an XF8 Analyzer (Seahorse Bioscience, Billerica, MA, USA). Human γδ T cells were adhered to the bottom of wells using Cell-Tak (#10317081, Fisher Scientific, Hampton, NH, USA) and incubated for 45 minutes at 37°C without CO2 before starting the assay. Specific mitochondrial respiration was determined after specific (BrHPP) or non-specific (PMA/ionomycin) activation.

### Statistical Analysis

Data were analyzed using GraphPad Prism 6.0 software (GraphPad Software, San Diego, CA, USA) and expressed as mean ± SD. Kruskal-Wallis and paired or unpaired non-parametric Student’s t-tests were used to determine significant differences. Significance was indicated as follows: *p < 0.05; **p < 0.01; ***p < 0.001.

## RESULTS

### Human γδ T cells from peripheral blood exhibit a Th1-biased functional profile

γδ T cells represented approximately 5% of lymphocytes in the peripheral blood of healthy adult donors (Figure 1A) and were isolated by negative selection for the subsequent experiments. After sorting, the recovered population was highly enriched in γδ T cells, with the majority of cells belonging to the Vδ2 positive compartment, although substantial inter-donor variability in the ratio of Vδ2+/V Vδ2-cells was observed (Figure 1A). To characterize their functional profile, cytokine secretion was analyzed using the LEGENDplex Human Th Cytokine Panel, which simultaneously quantifies 13 cytokines in the same supernatant. In the absence of stimulation, only IL-2 was consistently detected, whereas IFN-γ and TNF-α remained at/or near the detection threshold (2 pg/mL) (Figure 1B). Following strong, non-specific activation with PMA/ionomycin, IFN-γ and TNF-α became the dominant cytokines detected, while IL-2 increased only modestly, indicating a predominantly Th1-like response (Figure 1B). To determine which γδ T-cell subsets contributed to this response, intracellular IFN-γ staining was performed after PMA/ionomycin stimulation and compared between Vδ2+ and Vδ2-cells. Both subsets produced IFN-γ, but the response was markedly stronger in Vδ2+ cells, indicating that Vγ9Vδ2 T cells have a higher effector potential than Vδ2-γδ T cells (Figure 1C).

**Figure 1.**
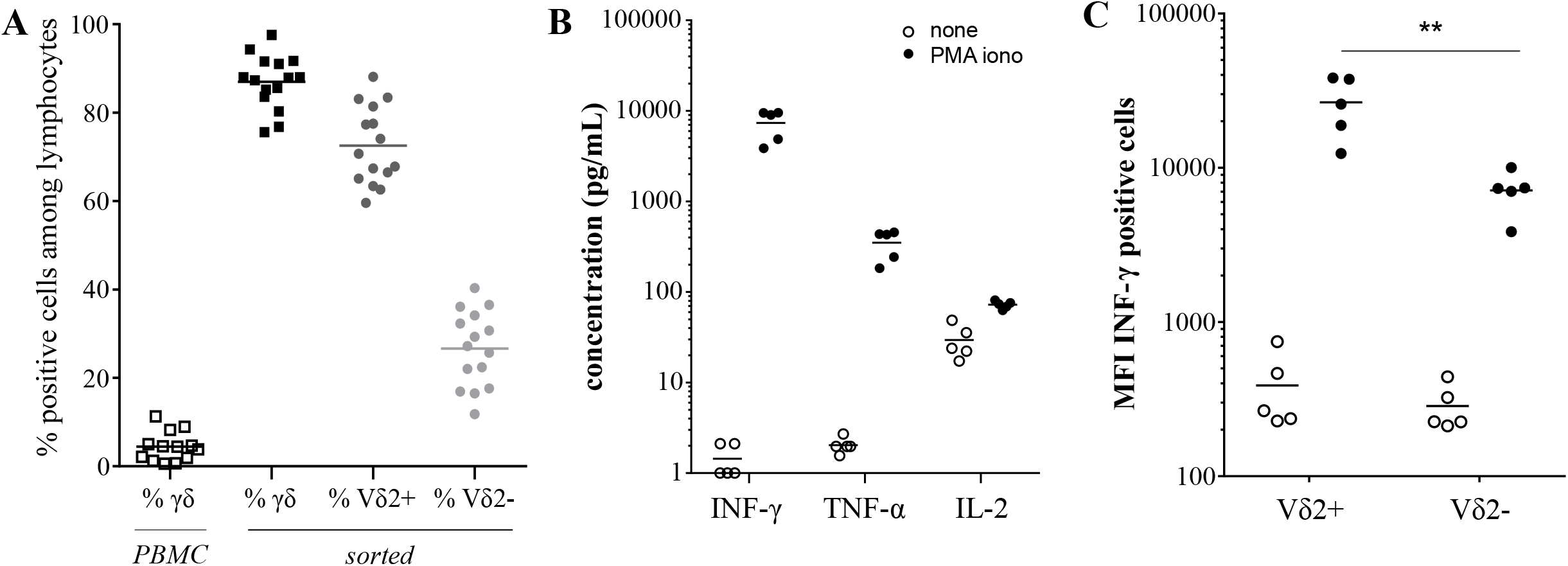
Peripheral blood γδ T cells display a pro-inflammatory functional profile. (A) Frequency of γδ T cells among PBMCs and distribution of Vδ2^+^ and Vδ2^−^ subsets after untouched sorting (n = 15 donors). (B) Cytokine secretion by sorted γδ T cells after 6 h stimulation with PMA/ionomycin or left unstimulated (none). Supernatants were analyzed using LEGENDplex Human Th Cytokine Panel. Only cytokines detected among the 13 assayed are shown (n = 5 donors). (C) Intracellular IFN-γ production after 6 h stimulation with PMA/ionomycin, analyzed in Vδ2^+^ and Vδ2^−^ subsets (n = 5 donors). ** p<0.01.

### Multiple amplification efficiently generates highly pure Vγ9Vδ2 T cells with an effector memory phenotype

Because Vγ9Vδ2 T cells are attractive candidates for cellular therapy due to their pleiotropic effector functions, we developed a multiple amplification protocol to obtain large numbers of highly pure cells from healthy donor blood (Figure 2A). After PBMC isolation, Vγ9Vδ2 T cells were specifically expanded using BrHPP, a synthetic phosphoantigen, or zoledronate, an aminobisphosphonate. Three weeks later, these cultures yielded resting polyclonal Vγ9Vδ2 T-cell populations with a purity of approximately 85% after BrHPP stimulation and 90% after zoledronate stimulation (Figure 2A-B). These cells were then re-stimulated non-specifically with PHA and feeder cells, and after three successive rounds of non-specific amplification, the resulting population remained highly enriched in Vγ9Vδ2 T cells, with a mean purity of 88% (Figure 2A-B). Subsequent analyses were performed at three defined stages of the protocol: untouched sorted γδ T cells from PBMCs (unT), specifically amplified Vγ9Vδ2 T cells (SA), and Vγ9Vδ2 T cells after three successive non-specific amplifications (3NSA). Phenotypic analysis further showed that the great majority of Vγ9Vδ2 T cells displayed an effector memory phenotype (CD27−CD45RA−), representing more than 90% of cells, and that this proportion significantly increased after each amplification step (Figure 2C).

**Figure 2.**
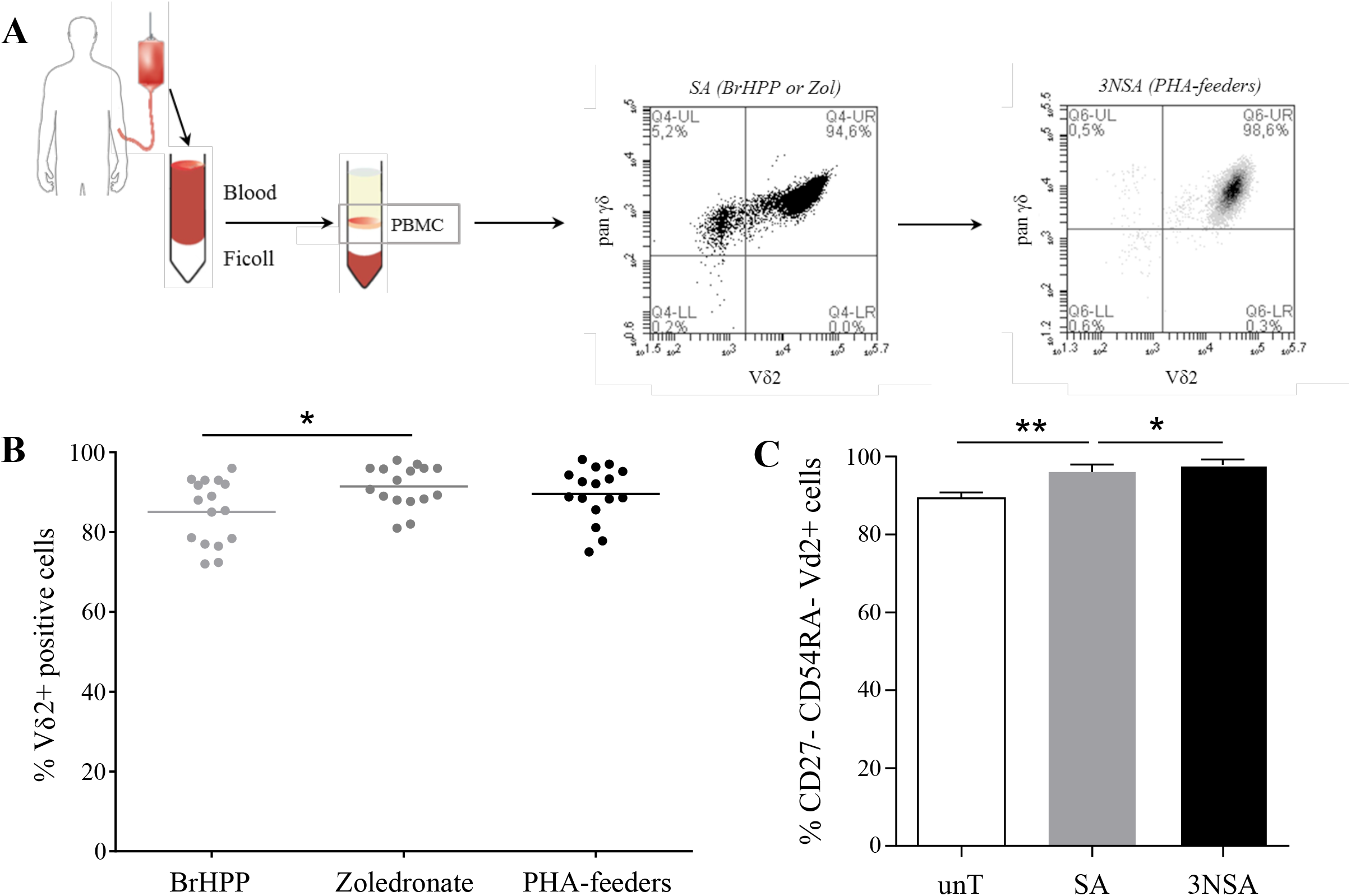
Stepwise amplification of Vγ9Vδ2 T cells preserves high purity and promotes differentiation. (A) Schematic representation of the amplification protocol. PBMCs were isolated from peripheral blood and Vγ9Vδ2 T cells were specifically expanded using BrHPP or Zoledronate (specific amplification, SA). After 3 weeks, cells returned to a resting state and were subsequently expanded through repeated non-specific stimulations using PHA and feeder cells (three rounds, 3NSA). Representative flow cytometry plots showing purity after SA (middle) and 3NSA (right) are displayed. (B) Quantification of Vδ2+ cell purity following specific amplification (BrHPP or zoledronate) and after three rounds of non-specific amplification (n = 16 donors). (C) Phenotypic characterization of differentiation status based on CD27 and CD45RA expression in untouched (unT), specifically amplified (SA), and three-times non-specifically amplified (3NSA) Vγ9Vδ2 T cells (n = 6 donors) * p<0.05 ; ** p<0.01.

### Amplified Vγ9Vδ2 T cells display enhanced effector functions together with reduced inhibitory checkpoint expression

We next assessed the functional consequences of amplification on Vγ9Vδ2 T cells. After 6 h of PMA/ionomycin stimulation, the same three cytokines identified in freshly isolated γδ T cells—IL-2, IFN-γ, and TNF-α—were detected, but their relative levels changed according to the amplification stage. Specific amplification and repeated non-specific amplification markedly increased IFN-γ and TNF-α secretion, while progressively decreasing IL-2 production, indicating reinforcement of a pro-inflammatory effector program (Figure 3A). This shift was confirmed by intracellular IFN-γ staining, which showed stronger IFN-γ expression after amplification, with no major difference between SA and 3NSA cells (Figure 3B). The anti-tumor reactivity of Vγ9Vδ2 T cells was then evaluated using PC3 prostate cancer cells sensitized with increasing concentrations of zoledronate. Both SA and 3NSA cells displayed significantly higher CD107a expression than untouched γδ T cells, although no relevant difference was observed between the two amplified conditions, indicating that the gain in cytotoxic responsiveness was already achieved after the first amplification step and maintained thereafter (Figure 3C). This functional enhancement was associated with a marked reduction in inhibitory checkpoint expression, with strong decreases in PD-1 and LAG-3, near-complete loss of CTLA-4, and no substantial change in TIM-3 expression (Figure 3D).

**Figure 3.**
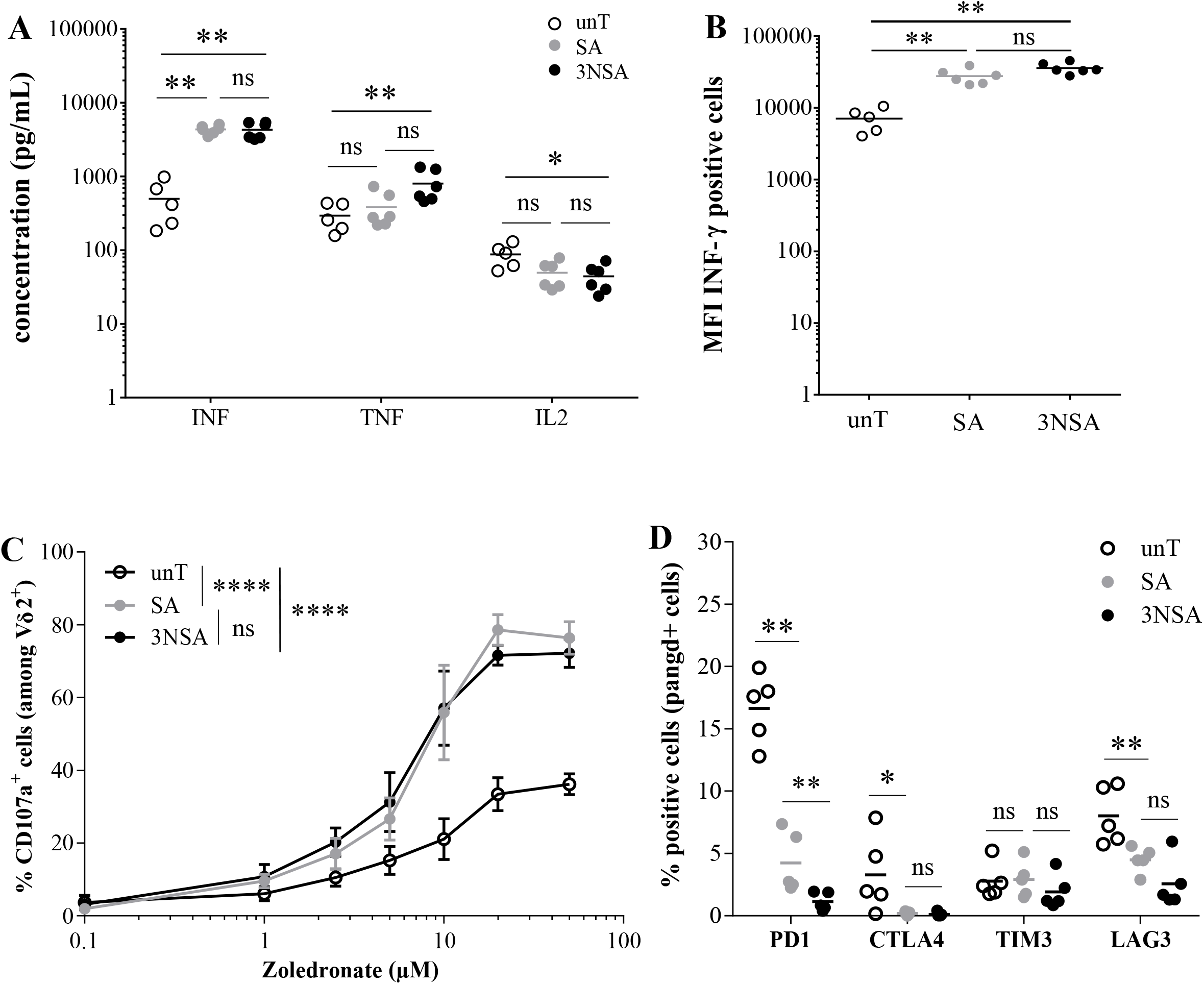
Functional competence of Vγ9Vδ2 T cells is enhanced after amplification. (A) Cytokine secretion following 6 h PMA/ionomycin stimulation in untouched (unT), specifically amplified (SA), and three-times non-specifically amplified (3NSA) Vγ9Vδ2 T cells, measured using LEGENDplex (n = 5 donors). (B) Intracellular IFN-γ production assessed after 6 h stimulation (n = 5 donors). (C) Degranulation capacity evaluated by CD107a expression after co-culture with zoledronate-sensitized PC3 prostate cancer cells (E:T ratio 1:1, 4 h).Dose-response curves are shown (n = 6 experiments from 3 donors). (D) Expression of inhibitory immune checkpoints (PD-1, CTLA-4, TIM-3, LAG-3) analyzed by flow cytometry (n = 5 donors) * p<0.05 ; ** p<0.01 ; **** p<0.0001.

### Amplification is associated with metabolic priming and progressive glycolytic reprogramming

We then investigated whether the functional changes induced by amplification were accompanied by metabolic remodeling. Seahorse analysis showed that BrHPP stimulation induced only a modest increase in oxygen consumption rate (OCR) and extracellular acidification rate (ECAR), whereas PMA/ionomycin triggered a stronger metabolic response followed by a progressive return toward baseline, except in 3NSA cells, which maintained higher activity over time (Figure 4A-B). At baseline, amplified Vγ9Vδ2 T cells displayed significantly higher metabolic activity than untouched γδ T cells, with SA cells showing increased ECAR and 3NSA cells showing increases in both OCR and ECAR, consistent with a metabolically primed effector memory state (Figure 4C). Analysis of the OCR/ECAR ratio further indicated that untouched γδ T cells relied predominantly on oxidative metabolism, whereas amplification progressively shifted Vγ9Vδ2 T cells toward a more glycolytic profile, particularly after repeated non-specific amplification (Figure 4D). These differences were most evident after BrHPP stimulation, while they were less pronounced after strong PMA/ionomycin stimulation, especially for OCR changes (Figure 4E-F). Together, these results indicate that amplification not only enhances effector functions, but also promotes metabolic reprogramming compatible with faster reactivation capacity, which may influence persistence and efficacy after adoptive cell transfer.

**Figure 4.**
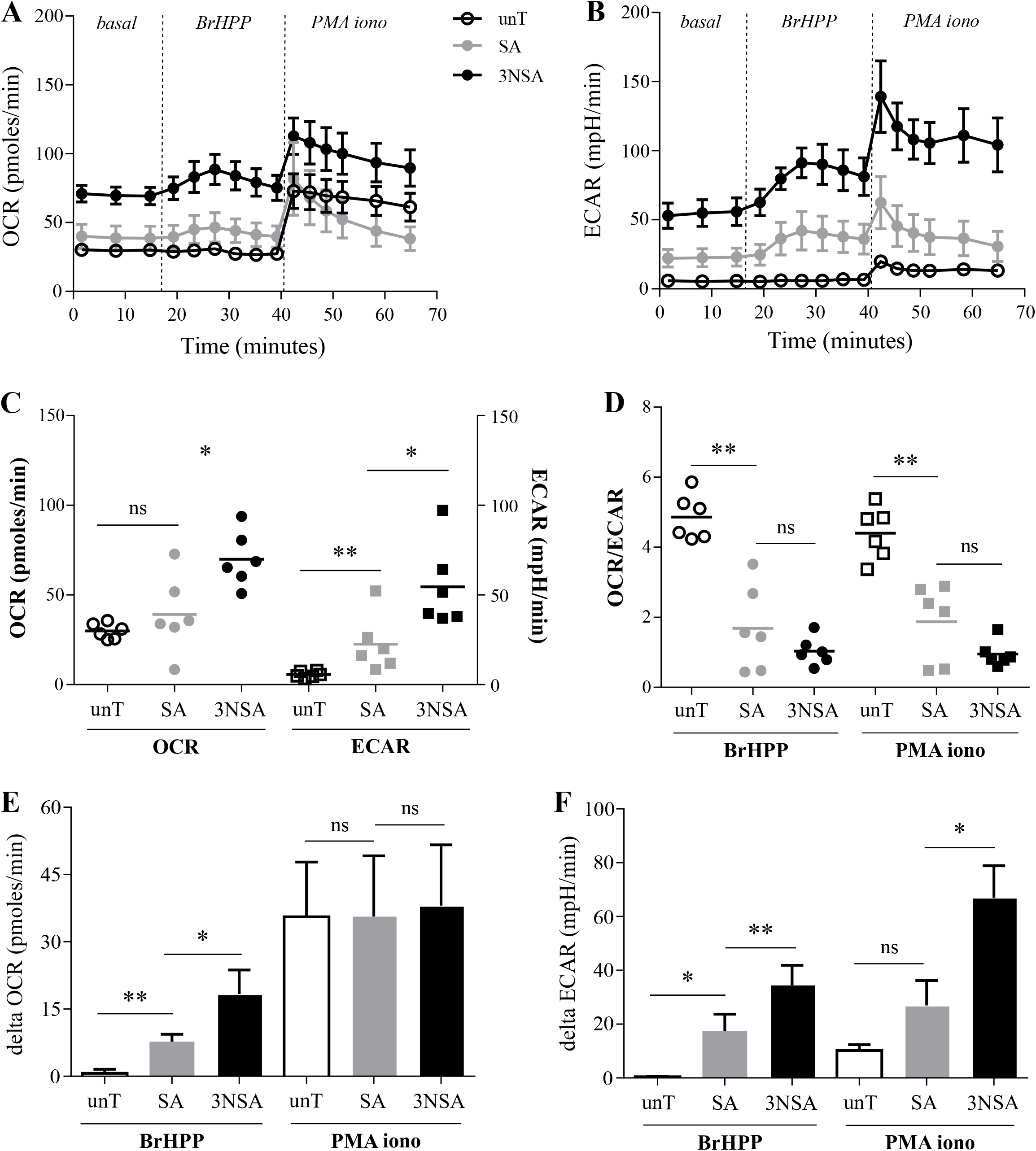
Metabolic reprogramming of Vγ9Vδ2 T cells during amplification. (A–B) Representative Seahorse profiles of oxygen consumption rate (OCR, A) and extracellular acidification rate (ECAR, B) at baseline, after specific activation (BrHPP), and after non-specific activation (PMA/ionomycin). (C) Basal OCR and ECAR values (n = 6 measurements from 3 donors). (D) OCR/ECAR ratio following BrHPP or PMA/ionomycin stimulation. (E–F) Metabolic response quantified as the difference (Δ) between baseline and post-stimulation values for OCR (E) and ECAR (F) (n = 6 measurements from 3 donors) * p<0.05 ; ** p<0.01.

### Repeated non-specific amplification enables extensive expansion while largely preserving purity and reactivity

To assess the suitability of this strategy for generating allogeneic Vγ9Vδ2 T-cell banks, we extended the non-specific amplification protocol until exhaustion and evaluated proliferation capacity, purity, and functional reactivity after each round of PHA-feeder stimulation. From the first to the fifth non-specific amplification, Vγ9Vδ2 T cells underwent approximately nine divisions per round, whereas a marked reduction in proliferation was observed only after the sixth stimulation (Figure 5A). Based on these results, we estimate that Vγ9Vδ2 T cells can undergo roughly 60 divisions from blood isolation to exhaustion, indicating a substantial expansion potential. Importantly, purity remained high throughout repeated amplification, with only a moderate decline at late stages (Figure 5B), and anti-tumor reactivity, assessed by CD107a expression, was largely maintained over successive stimulation cycles, although a reduction became apparent after the fifth non-specific amplification (Figure 5C). Altogether, these data indicate that repeated non-specific amplification allows large-scale production of Vγ9Vδ2 T cells while preserving the main characteristics required for therapeutic use.

**Figure 5.**
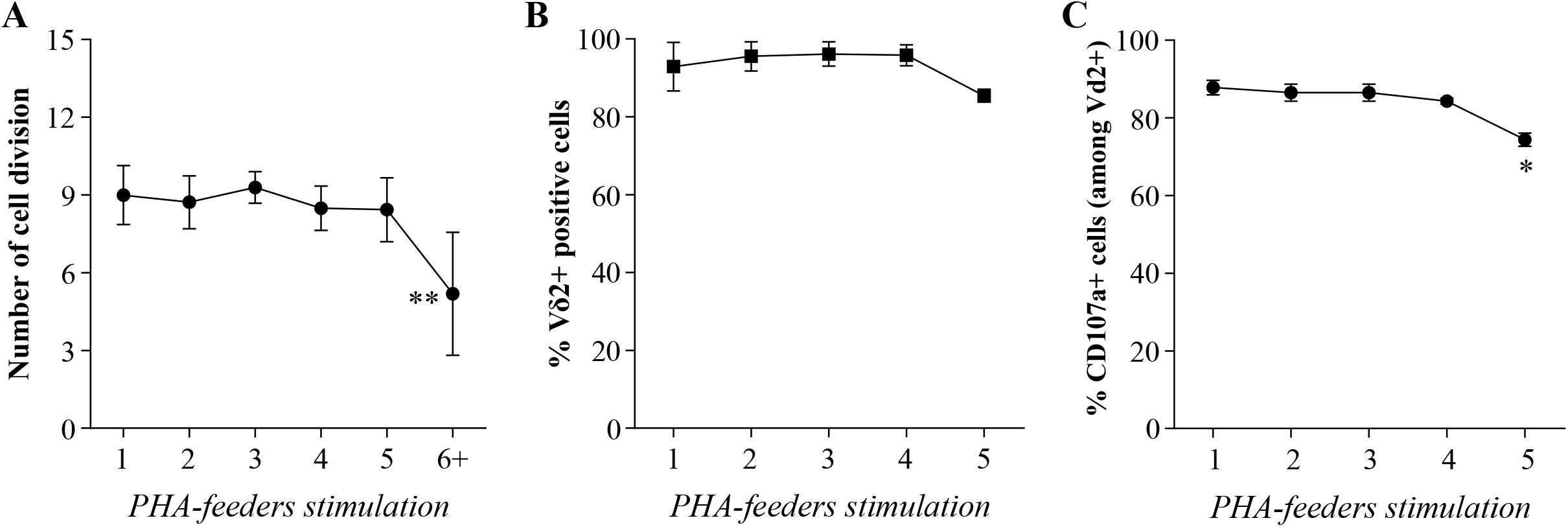
Repeated non-specific stimulation poorly affect purity and reactivity. (A) Number of cell divisions after successive rounds of PHA/feeder stimulation. (B) Maintenance of Vδ2+ cell purity across amplification cycles. (C) Functional reactivity assessed by CD107a expression following each stimulation round (n = 3–6 per condition) * p<0.05 ; ** p<0.01.

## DISCUSSION

Vγ9Vδ2 T cells are attractive candidates for cancer immunotherapy because they combine broad antitumor reactivity, major histocompatibility complex-independent recognition, and low alloreactivity, making them well suited for allogeneic adoptive cell transfer strategies (Wilhelm et al., 2003; Dieli et al., 2007; Silva-Santos et al., 2015). In the present study, we show that successive rounds of specific and non-specific ex vivo amplification generate highly enriched polyclonal Vγ9Vδ2 T-cell populations while preserving, and in some respects enhancing, their functional properties. These data support the feasibility of producing large numbers of Vγ9Vδ2 T cells for off-the-shelf therapeutic applications.

Our results first confirm that peripheral blood γδ T cells from healthy donors display a predominantly pro-inflammatory functional profile, characterized by strong IFN-γ and TNF-α production upon activation, with a greater contribution from the Vδ2+ compartment. Importantly, this profile was maintained throughout the amplification process. On the contrary, repeated amplification was associated with increased IFN-γ and TNF-α secretion together with reduced IL-2 production, consistent with reinforcement of an effector program. This was accompanied by sustained degranulation against zoledronate-sensitized tumor cells, indicating that repeated expansion does not compromise antitumor reactivity.

A major challenge for γδ T-cell-based immunotherapy is the development of scalable manufacturing strategies capable of generating standardized cell products. Here, the combination of an initial specific expansion step using BrHPP or zoledronate with repeated non-specific restimulation using PHA and feeder cells resulted in highly pure Vγ9Vδ2 T-cell populations that remained enriched after several rounds of amplification. In parallel, amplified cells displayed a predominantly effector memory phenotype, which became even more pronounced during expansion. These features are particularly relevant for adoptive cell therapy, as they suggest that large-scale production can be achieved without major loss of cellular identity or immediate effector potential.

Another notable finding is the reduction in inhibitory immune checkpoint expression after amplification. PD-1, CTLA-4, and LAG-3 were all decreased, whereas TIM-3 remained relatively stable. This phenotype is consistent with the enhanced functional responses observed in amplified cells. Although checkpoint expression measured ex vivo cannot predict how these cells will behave after transfer into an immunosuppressive tumor microenvironment, the low expression of several inhibitory receptors nevertheless represents a favorable characteristic for therapeutic use.

Our study also shows that repeated amplification is accompanied by marked metabolic remodeling. Amplified Vγ9Vδ2 T cells exhibited increased basal metabolic activity compared with freshly isolated γδ T cells, and repeated stimulation progressively shifted them toward a more glycolytic profile. This pattern is compatible with acquisition of an effector memory-like state and may contribute to the faster reactivation capacity observed after stimulation. Given the growing importance of metabolic fitness in adoptive cell therapy, these findings suggest that metabolic parameters should be considered alongside phenotype and function when optimizing γδ T-cell manufacturing protocols.

These observations are in line with previous preclinical and clinical studies supporting the therapeutic potential of Vγ9Vδ2 T cells. In particular, zoledronate-based approaches have shown that increasing intracellular phosphoantigen levels in tumor cells enhances their recognition by Vγ9Vδ2 T cells (Gober et al., 2003). Our results extend this rationale by indicating that repeated amplification can increase cell yield without immediate functional loss, which may help overcome one of the main practical limitations of current γδ T-cell approaches.

This study has several limitations. First, all experiments were performed with cells from healthy donors, and the behavior of patient-derived Vγ9Vδ2 T cells may differ. Second, antitumor activity was mainly assessed in vitro through degranulation assays and will require confirmation in vivo. Third, although functionality was maintained over several rounds of amplification, proliferation eventually declined, indicating that expansion capacity is substantial but not unlimited. Defining the optimal balance between cell yield and late-stage functional exhaustion will therefore be important for clinical translation.

Overall, our data show that successive specific and non-specific amplification steps can generate highly pure, polyclonal Vγ9Vδ2 T-cell populations with preserved effector functions, reduced expression of several inhibitory checkpoints, and substantial proliferative potential. These findings strengthen the rationale for developing Vγ9Vδ2 T cells as allogeneic, off-the-shelf products for adoptive immunotherapy and provide a framework for further optimization of γδ T-cell manufacturing strategies.

## Acknowledgements

The authors acknowledge the Cytocell - Flow Cytometry and FACS core facilty (SFR Bonamy, BioCore, Inserm UMS 016, CNRS UAR 3556, Nantes, France) for its technical expertise and help. The authors also acknowledge Ophélie Renoult and Claire Pecqueur (CRCINA, INSERM, CNRS, Université d’Angers, Université de Nantes, Nantes, France) for help to performed seahorse experiments. This project was funded by the French National Institute of Health and Medical Research (INSERM), providing the infrastructure and resources necessary to conduct this research.

## REFERENCES

Bennouna J, Bompas E, Neidhardt EM, Rolland F, Philip I, Galéa C, Salot S, Saiagh S, Audrain M, Rimbert M et al. 2008 Phase-I study of Innacell gammadelta, an autologous cell-therapy product highly enriched in gamma9delta2 T lymphocytes, in combination with IL-2, in patients with metastatic renal cell carcinoma. Cancer Immunology, Immunotherapy: CII 57 1599–1609. (doi:10.1007/s00262-008-0491-8)

Cano CE, Pasero C, De Gassart A, Kerneur C, Gabriac M, Fullana M, Granarolo E, Hoet R, Scotet E, Rafia C et al. 2021 BTN2A1, an immune checkpoint targeting Vγ9Vδ2 T cell cytotoxicity against malignant cells. Cell Reports 36 109359. (doi:10.1016/j.celrep.2021.109359)

Chauvin C, Joalland N, Perroteau J, Jarry U, Lafrance L, Willem C, Retière C, Oliver L, Gratas C, Gautreau-Rolland L et al. 2019 NKG2D Controls Natural Reactivity of Vγ9Vδ2 T Lymphocytes against Mesenchymal Glioblastoma Cells. Clinical Cancer Research: An Official Journal of the American Association for Cancer Research 25 7218–7228. (doi:10.1158/1078-0432.CCR-19-0375)

Chien Y, Meyer C & Bonneville M 2014 γδ T cells: first line of defense and beyond. Annual Review of Immunology 32 121–155. (doi:10.1146/annurev-immunol-032713-120216)

Cimmino L, Dolgalev I, Wang Y, Yoshimi A, Martin GH, Wang J, Ng V, Xia B, Witkowski MT, Mitchell-Flack M et al. 2017 Restoration of TET2 Function Blocks Aberrant Self-Renewal and Leukemia Progression. Cell 170 1079-1095.e20. (doi:10.1016/j.cell.2017.07.032)

Dieli F, Vermijlen D, Fulfaro F, Caccamo N, Meraviglia S, Cicero G, Roberts A, Buccheri S, D’Asaro M, Gebbia N et al. 2007 Targeting human {gamma}delta} T cells with zoledronate and interleukin-2 for immunotherapy of hormone-refractory prostate cancer. Cancer Research 67 7450–7457. (doi:10.1158/0008-5472.CAN-07-0199)

Dunne MR, Mangan BA, Madrigal-Estebas L & Doherty DG 2010 Preferential Th1 cytokine profile of phosphoantigen-stimulated human Vγ9Vδ2 T cells. Mediators of Inflammation 2010 704941. (doi:10.1155/2010/704941)

Fournié J-J, Sicard H, Poupot M, Bezombes C, Blanc A, Romagné F, Ysebaert L & Laurent G 2013 What lessons can be learned from γδ T cell-based cancer immunotherapy trials? Cellular & Molecular Immunology 10 35–41. (doi:10.1038/cmi.2012.39)

Gober H-J, Kistowska M, Angman L, Jenö P, Mori L & De Libero G 2003 Human T Cell Receptor γδ Cells Recognize Endogenous Mevalonate Metabolites in Tumor Cells. The Journal of Experimental Medicine 197 163–168. (doi:10.1084/jem.20021500)

Harly C, Guillaume Y, Nedellec S, Peigné C-M, Mönkkönen H, Mönkkönen J, Li J, Kuball J, Adams EJ, Netzer S et al. 2012 Key implication of CD277/butyrophilin-3 (BTN3A) in cellular stress sensing by a major human γδ T-cell subset. Blood 120 2269–2279. (doi:10.1182/blood-2012-05-430470)

Jarry U, Chauvin C, Joalland N, Léger A, Minault S, Robard M, Bonneville M, Oliver L, Vallette FM, Vié H et al. 2016 Stereotaxic administrations of allogeneic human Vγ9Vδ2 T cells efficiently control the development of human glioblastoma brain tumors. OncoImmunology 5 e1168554. (doi:10.1080/2162402X.2016.1168554)

Joalland N & Scotet E 2020 Emerging Challenges of Preclinical Models of Anti-tumor Immunotherapeutic Strategies Utilizing Vγ9Vδ2 T Cells. Frontiers in Immunology 11. (doi:10.3389/fimmu.2020.00992)

Joalland N, Chauvin C, Oliver L, Vallette FM, Pecqueur C, Jarry U & Scotet E 2018 IL-21 Increases the Reactivity of Allogeneic Human Vγ9Vδ2 T Cells Against Primary Glioblastoma Tumors. Journal of Immunotherapy (Hagerstown, Md.: 1997) 41 224–231. (doi:10.1097/CJI.0000000000000225)

Joalland N, Lafrance L, Oullier T, Marionneau-Lambot S, Loussouarn D, Jarry U & Scotet E 2019 Combined chemotherapy and allogeneic human Vγ9Vδ2 T lymphocyte-immunotherapies efficiently control the development of human epithelial ovarian cancer cells in vivo. OncoImmunology 8 e1649971. (doi:10.1080/2162402X.2019.1649971)

Rochman Y, Spolski R & Leonard WJ 2009 New insights into the regulation of T cells by gamma(c) family cytokines. Nature Reviews. Immunology 9 480–490. (doi:10.1038/nri2580)

Rosenberg SA & Restifo NP 2015 Adoptive cell transfer as personalized immunotherapy for human cancer. Science (New York, N.Y.) 348 62–68. (doi:10.1126/science.aaa4967)

Santolaria T, Robard M, Léger A, Catros V, Bonneville M & Scotet E 2013 Repeated systemic administrations of both aminobisphosphonates and human Vγ9Vδ2 T cells efficiently control tumor development in vivo. Journal of Immunology (Baltimore, Md.: 1950) 191 1993–2000. (doi:10.4049/jimmunol.1300255)

Silva-Santos B, Serre K & Norell H 2015 γδ T cells in cancer. Nature Reviews. Immunology 15 683–691. (doi:10.1038/nri3904)

Tanaka Y, Morita CT, Tanaka Y, Nieves E, Brenner MB & Bloom BR 1995 Natural and synthetic non-peptide antigens recognized by human gamma delta T cells. Nature 375 155–158. (doi:10.1038/375155a0)

Vantourout P & Hayday A 2013 Six-of-the-best: unique contributions of γδ T cells to immunology. Nature Reviews. Immunology 13 88–100. (doi:10.1038/nri3384)

Wilhelm M, Kunzmann V, Eckstein S, Reimer P, Weissinger F, Ruediger T & Tony H-P 2003 Gammadelta T cells for immune therapy of patients with lymphoid malignancies. Blood 102 200–206. (doi:10.1182/blood-2002-12-3665)

Zhang P, Zhang G & Wan X 2023 Challenges and new technologies in adoptive cell therapy. Journal of Hematology & Oncology 16 97. (doi:10.1186/s13045-023-01492-8)

Zhao Y, Niu C & Cui J 2018 Gamma-delta (γδ) T cells: friend or foe in cancer development? Journal of Translational Medicine 16 3. (doi:10.1186/s12967-017-1378-2)

